# Seasonal fluctuations in the bone microstructure of *Sciurus vulgaris fuscoater* humeri: a case study using phenomics on µCT-scans

**DOI:** 10.1101/2023.12.10.571007

**Authors:** AH van Heteren, AS Luft, M Toth, J Dewanckele, M Marsh, J De Beenhouwer

## Abstract

*Sciurus vulgaris* Linnaeus, 1758, the red squirrel, is a small, mostly arboreally living rodent, spread across the Palearctic. It is mostly vegetarian, feeding on plants, fungi and seeds, and is less active in the winter months, but does not hibernate. In this lateral study, the humeri of the subspecies *Sciurus vulgaris fuscoater*, the Central European red squirrel, were analysed to uncover potential intraspecific variation between individuals found in different seasons.

The µCT-scans were obtained with a resolution of 26 microns. Five bone parameters were calculated and statistically evaluated with regards to seasonal variations: total volume, bone volume, endocortical surface, cortical thickness, and average trabecular thickness.

Bone volume, trabecular thickness and endocortical thickness correlate with bone size, whereas cortical thickness does not. Seasonal differences were observed between the warmer summer and autumn months versus the colder winter and spring months for all parameters. We, speculatively, relate the observed seasonal variation to nutrient intake, notably calcium. These results offer a deeper understanding of intraindividual variation in red squirrels, that may be useful in further ecological, taxonomic, and paleontological research.

## Introduction

The importance of seasonality is undeniable. Yearly temperature and precipitation cycles are fundamental to the availability of food and water to animals (Kwiecien *et al*. 2022). Due to global climate change, seasonality is changing in many parts of the world (Marelle *et al*. 2018; Santer *et al*. 2018). It is important to understand the physiological response of animals to seasonality to comprehend the challenges they might face soon. Seasonal adaptations in mammals are widely reported and mostly concern pelage and adiposity (e.g, Scherbarth & Steinlechner 2010; Zimova *et al*. 2018). There is limited research on the response of cortical bone to seasonality during growth (Köhler *et al*. 2012), but, until now, the response of trabecular bone has remained unknown. Although, the trabecular and cortical architecture of two squirrel femora has been studied (Mielke *et al*. 2018), such a small sample size does not allow for an analysis of seasonality. Here, we will analyse the trabecular bone structure of the red squirrel (*Sciurus vulgaris* Linnaeus, 1758) to assess whether this mammal displays a physiological response to seasonality.

The red squirrel is a medium-sized, arboreal mammal belonging to the order Rodentia, and the family Sciuridae (Lurz *et al*. 2005). The red squirrel is one of the most common squirrel species worldwide, populating big parts of the European and Asian continents. Living an arboreal lifestyle, their preferred habitat is the deciduous and coniferous forest, where they build their nests, also called dreys. More than 40 subspecies have been described, not all of them valid (Corbet 1978). It has previously been observed, that *S. vulgaris fuscoater* has a larger geographic variation in skull size, than other subspecies (Wiltafsky 1978). Red squirrels are found in forest regions across the Palearctic from the Iberian Peninsula and Britain, all the way to Japan (Lee & Fukuda 1999; Thorington Jr & Hoffmann 2005). They are found mainly in conifer forests or mixed woodland to provide for a year-round sufficient diet (Moller 1983a; Moller 1983b; Lurz *et al*. 1995; Lurz *et al*. 1998).

Red squirrels have a wide-ranging diet, consisting mainly of the fruits and seeds of different tree species but can also include eggs and small birds, depending on seasonal availability (Lurz *et al*. 2005). Overall, foraging behaviour in red squirrels is highly dependent on their environment (Krauze-Gryz & Gryz 2015). Squirrel body mass does not increase in autumn in preparation for winter (A. Wauters *et al*. 2007) and, in the winter, red squirrels do not go into hibernation, but they adapt their foraging behaviour, looking for more high-energy resources, such as pine seeds, (Krauze-Gryz & Gryz 2015). Their most important food sources are conifer seeds, fungi, nuts, fruits, buds, and catkins (Moller 1983a; Moller 1983b), but knowledge on their scatter-hoarding behaviour and the influence on diet composition throughout the year is still very limited (Krauze-Gryz & Gryz 2015) and mostly anecdotal. A study from Poland showed that, when offered supplemental feeding, 80% of the animals took supplemental nuts in winter, but only 67% took supplemental nuts in autumn (Kostrzewa & Krauze-Gryz 2020), suggesting that, despite scatter-hoarding, nuts are less readily available in winter than in autumn. Tree buds and flowers were a significant part of the red squirrel diet (>70%) during late winter and spring in England (Shuttleworth 1997). The only study, to our knowledge, that is based on stomach contents comes from East Scotland and only lists occurrence data (Tittensor 1970).

Bone structure and composition can provide information about taxonomic affiliation, age, health status, and life history, thus making it an important study material in biology (Boskey & Coleman 2010; Barak *et al*. 2013a; Barak *et al*. 2013b; Meier *et al*. 2013; Amson *et al*. 2017; de Bakker *et al*. 2018).

Bone parameters can be quantified and analysed, enabling conclusions to be drawn about the organism and its life circumstances (Mullender *et al*. 1996; Doube *et al*. 2011; Chirchir *et al*. 2017; Tsegai *et al*. 2018). As bone mineralization and bone microstructure are dependent on nutrient intake (Scholz-Ahrens & Schrezenmeir 2007; De Cuyper *et al*. 2020), a fluctuating food availability across seasons could result in detectable changes in squirrel bones.

While intraspecific microstructure variation has been studied in human skeletons (e.g., Saers *et al*. 2018; Vom Scheidt *et al*. 2019), the focus in other animals has been mainly on interspecific variation, caused by, for example, different forms of locomotion, adaptation to their environment and evolutionary history (e.g., Meier *et al*. 2013; Mielke *et al*. 2018). Studies focusing on interspecific variation in bone microstructure use a relatively low sample size for each species (Barak *et al*. 2013a; Meier *et al*. 2013; Amson *et al*. 2017; Mielke *et al*. 2018), even though it is not necessarily true that the individuals chosen are representative for the entire population or species. To determine the extent of intraspecific variation, for example associated with seasonality, a large sample of specimens belonging to the same species, in this case *S. vulgaris fuscoater*, needs to be quantified, analysed, and statistically tested.

## Materials

The specimens used in this study are listed in Suppl. Info 1. All specimens belong to the subspecies *Sciurus vulgaris fuscoater* and are from Bavaria (Germany). They entered the Bavarian State Collection of Zoology between April 1907 and February 1917, and they consist of complete skeletons. It is not known how the specimens were collected but given their completeness, it seems likely that they were either trapped or hunted, rather than coincident finds of dead animals.

A total of 40 humeral bones, belonging to mature individual based on the external morphology of the skeletons and absence of a symphysial line, were scanned. The humerus was chosen, because previous research on dogs suggests that the proximal-most bones of weight-bearing limbs show the smallest anabolic and catabolic responses to exercise and disuse, respectively (Jaworski *et al*. 1980; Turner 1999; Robling *et al*. 2006), possibly because the proximal-most bones are loaded more indirectly (Robling *et al*. 2006). Furthermore, empirical studies on other sciurids showed that the humerus developed significantly lower stresses than the radius and the ulna (Biewener 1983). Additionally, the humerus is expected to experience less substrate reaction forces than the femur (Andrada *et al*. 2013), the most proximal bone in the hind limb. As such, it would logically follow that the humerus would be less influenced by load and could respond more freely to environmental factors, such as seasonal changes. For the trabecular analyses, the proximal trabeculae were chosen, because trabecular morphology has a functional significance. For example, once a discontinuity in a trabecular element is created, that element can no longer support load (Nazarian *et al*. 2008). As the cross-struts between longitudinally oriented trabeculae become disconnected, the remaining trabeculae become functionally longer and weaker. Previous research in dogs, however, suggests that the proximal-most bones of weight-bearing limbs are loaded more indirectly, and that interstitial fluid pressure could be important for bone maintenance (Jaworski *et al*. 1980; Turner 1999; Robling *et al*. 2006). As such, it would logically follow that the proximal part of the humerus could respond to environmental factors, such as seasonal changes, without impeding functionality.

Collection year, month, and day (presumably within days of death) were available for all bones except one, which was only labelled January 1915. For this specimen, we used the 15^th^ of January, as it is the middle of the month, when a more precise date was needed in the analyses. The weather from 1907 to 1917 was comparable to preceding and following decades (meteo.plus 2023). A multivariate multiple linear regression shows there is no relationship between bone anatomy and annual temperature or annual precipitation for that period (Overall multivariate analysis of variance, Wilks’ lambda=0.7098, F(10, 66)=1.234, p=0.2864, with each of the regression coefficients p>0.0669, R^2^<0.1131). Therefore, weather fluctuations over this 11-year period should not influence the analyses. As might be expected, however, the temperatures the squirrels in this study (11-year period between 1907 and 1917) and more recent squirrels (11-year period between 2012 and 2022) were exposed to were significantly (t=7.3268, p<0.0001), but precipitation was similar (t=0.2384, p=0.8140) (data from meteo.plus (2023)). The effect of this temperature increase on weather fluctuations and squirrel behaviour is not yet clear at present.

Trailing and leading forelimbs serve different functions in red squirrels; the trailing forelimb functions as a shock absorber and the leading forelimb stabilises and supports the body (Schmidt 2011). Since it is impossible to know which forelimb was preferentially used in which function by the squirrels in this study, using only one side could bias the results. Therefore, where possible, both left and right bones of the same individual were included. Inclusion of bilateral data, however, assumes independence between paired data when in fact there might be dependence, increasing the likelihood of Type I error (Sullivan *et al*. 2016; Ying *et al*. 2018). On the other hand, a single measure per individual or an average of the paired measures is unnecessarily conservative and increases the likelihood of a Type II error (Camarillo *et al*. 2023). No significant difference was found between left and right bones in our dataset (Multivariate analysis of variance, Pillai trace=0.0708, F(5, 34)=0.5180, p=0.7608), so this should not have biased the relationships between the variables. For completeness, we have additionally done all statistical analyses presented below using the left-right means for those individuals for which both values were available (Suppl. Info. 2). The general patterns are the same, but the p-values are generally higher. These more conservative results do not change our interpretations.

The animals were grouped into four seasonal groups. Division into seasons was based on feeding behaviour (Moller 1983a) and the availability of tree seeds (Gurnell 1993): summer (June to September), fall (October and November), winter (December to February) and spring (March to May). It is worth noting that the present sample only contains 2 fall bones from the same individual. This is because red squirrels are difficult to catch in fall, when food is plentiful, and are often not sampled at all that time of year (e.g., Moller 1983a; Gurnell 1996). The two fall bones are not included in analyses with fall as a separate group, since the sample size would be smaller than k+1, where k is the number of groups.

## Methods

The µCT scans were obtained with a CoreTOM, located at Tescan in Ghent. The humeri were placed in individual plastic specimen jars and then stacked per 18 in a large PVC sample tube. A scan lasted 180 minutes per tube (10 minutes per humerus). Scans were taken at 160 kV, 0.156 mA and 25 W with 901 views over a 360° rotation per bone resulting in a 0.4° angular step size. The tube turned continuously rather than stepwise. The radiation source and detector were programmed to automatically move up to the next bone and take another 901 views. The reconstruction of the scans was largely automated using a Python script (Suppl. Info. 3; van Aarle *et al*. 2015; van Aarle *et al*. 2016) and the results were improved by removing ring artefacts. Each resultant image stack comprised of 2500 sectional images. The developmental stage of the squirrel (adult or subadult) was determined by the presence or absence of an epiphyseal plate in the trabecular bone of the proximal humerus.

The image stacks of the individual bones were read into Dragonfly 2021.1 with a voxel size of 0.026 mm. A python-based macro (Suppl. Info. 4) was used to create a 3D region of interest (ROI) that only includes the bone tissue of the humerus. The processing duration of the macro was approximately 5 min for each bone. The approximate thickness of the trabeculae was measured with the ruler tool for each individual, since the trabecular thickness varies intraspecifically. The Buie method, which is based on a dual threshold method, was then selected to start the automated segmentation between trabecular and cortical bone (Buie *et al*. 2007). This largely automatically generated segmentation required only minor manual corrections. Before calculating the parameters, the trabecular ROI was split by connected components and of the two largest connected components the proximal component was kept (*Figure 1*). This ensures all relevant trabeculae are included in the analysis and obviates the need to manually, and by human nature subjectively, choose a core-shaped (Benito *et al*. 2003; Benito *et al*. 2005), spherical (Skedros *et al*. 2012; Bachmann *et al*. 2022) or cubic (Hoechel *et al*. 2015; Amson *et al*. 2017; Marcián *et al*. 2017) ROI. After calculation of the parameters in the bone analysis module of the ORS Dragonfly software, the data was exported as CSV files and subjected to statistical testing.

**Figure 1:**
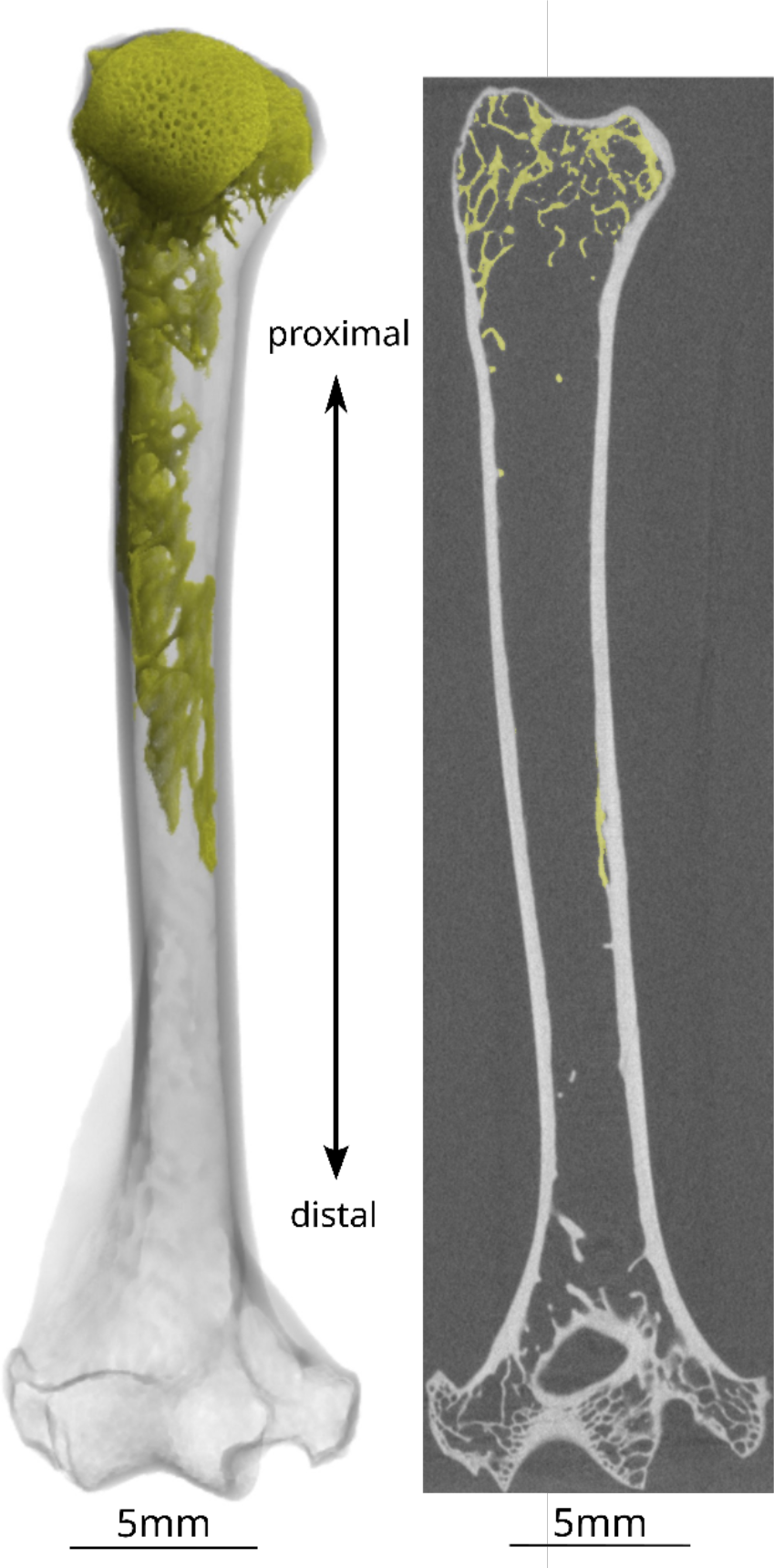
Example of the two largest connected components of trabecular bone (coloured) in a squirrel humerus. The region of interest for trabecular thickness in the proximal part is indicated in yellow. A. Longitudinal slice. B. 3D view with transparent cortical bone.

All statistical tests were conducted in the software Past 4.11 (Hammer *et al*. 2001) with a significance level of α = 0.05 and p-values that are below α = 0.01 are considered highly significant. All plots were made in Excel. The quartiles in the boxplots were calculated exclusive of the median and any datapoints beyond one and a half box lengths from either end of the box are considered outliers. The sinus trendlines in the scatterplots were created with Solver, a Microsoft Excel add-in. The regressions onto size were kept blind to season and season colour was only added later for illustrative purposes. In all other analyses (e.g., ANOVA) season or date was an integral part of the analysis. The following parameters were taken into consideration following the definitions of Bouxsein (Bouxsein *et al*. 2010), except endocortical surface (Object Research Systems 2019), but these parameters often have different abbreviations in the preclinical literature (Dempster *et al*. 2013):

Total volume (TV) = Volume of the entire region of interest (mm^3^), this includes non-bone spaces within the bone.

Bone volume (BV) = Volume of the region segmented as bone (mm^3^).

Endocortical surface (Ec.S3D) = Endocortical surface (mm^2^), assessed using direct 3D methods.

Average cortical thickness (Ct.Th.) = Mean cortical thickness (mm).

Trabecular thickness of the proximal trabeculae (Tb.Th.prox) = Mean thickness of trabeculae (mm), assessed using direct 3D methods, as applied to the largest continuous network of trabeculae in the proximal part of the bone.

Total volume would be a logical candidate for a measure of absolute size of the bone that includes both a length and a robusticity component. ANOVAs were performed on total volume to assure that it was not influenced by any seasonal variations (see below for how the assumptions were tested). The dependence of the variables of interest on size was tested using a multivariate regression of those variables onto total volume as a proxy for bone size. In those instances where the regression was significant, analyses were continued with the regression residuals. When the regression was non-significant, analyses were continued with the raw data.

Levene’s tests for homogeneity of variance from means and from medians were performed, as well as the Shapiro-Wilk test for normal distribution. Parametric testing was only continued if all these tests provided insignificant results, which was fortunately the case for all variables.

To test for any seasonal differences in the bone microstructure of squirrels, multivariate analyses of variance (MANOVAs) were performed for the estival (summer) semiyear vs the hibernal (winter) semiyear, as well as for the four seasons on cortical thickness, proximal trabecular thickness, endocortical surface and bone volume. Hotelling’s T^2^ analyses were conducted to determine where in the data the significance arises. The p-values and the Mahalanobis D^2^ effect size were reported.

To determine which factors might be important, individual analyses of variance (ANOVAs) were performed on cortical thickness, proximal trabecular thickness, endocortical surface and bone volume for seasonality, and in case of significant findings additional Tukey’s pairwise tests were performed. Since these are Model II (random effects) ANOVAs, the intraclass correlation coefficient (ICC) is also given for significant results, in addition to the customary parameters.

## Results

Total volume was found to be independent of estival or hibernal semiyear (F(1, 38)=2.111, p=0.1545) and of the four seasons (F(3, 36)=0.874, p=0.4638). As such, it can be used as an independent proxy for bone size to be used in the regression analyses.

Bone volume and endocortical surface were highly significantly correlated with total volume, but cortical thickness and proximal trabecular thickness were not (*Table 1* and *Figure 2*). Bone volume and endocortical surface show a significant correlation with total volume. For those variable, subsequent analyses were done on the regression residuals. For cortical thickness and trabecular thickness, the raw data were used.

**Table 1:** Results of the multivariate regression of bone volume (BV), cortical thickness (Ct.Th.), endocortical surface (Ec.S3D) and thickness of the proximal trabeculae (Tb.Th. prox) onto total volume. The significant values are indicated in bold font.

| *Variable* | *Slope* | *Error* | *Intercept* | *Error* | *r* | *p* |
| --- | --- | --- | --- | --- | --- | --- |
| *BV* | 0.39833 | 0.04679 | 45.422 | 22.589 | 0.80997 | **2.45E-10** |
| *Ec.S3D* | 0.87231 | 0.04303 | 104.500 | 21.258 | 0.95485 | **1.25E-21** |
| *Ct.Th.* | 4.23E-06 | 0.00010 | 0.421 | 0.049 | 0.00680 | 0.96677 |
| *Tb.Th. prox* | 5.73E-05 | 2.34E-05 | 0.091 | 0.011 | 0.36942 | **0,0190** |

**Figure 2:**
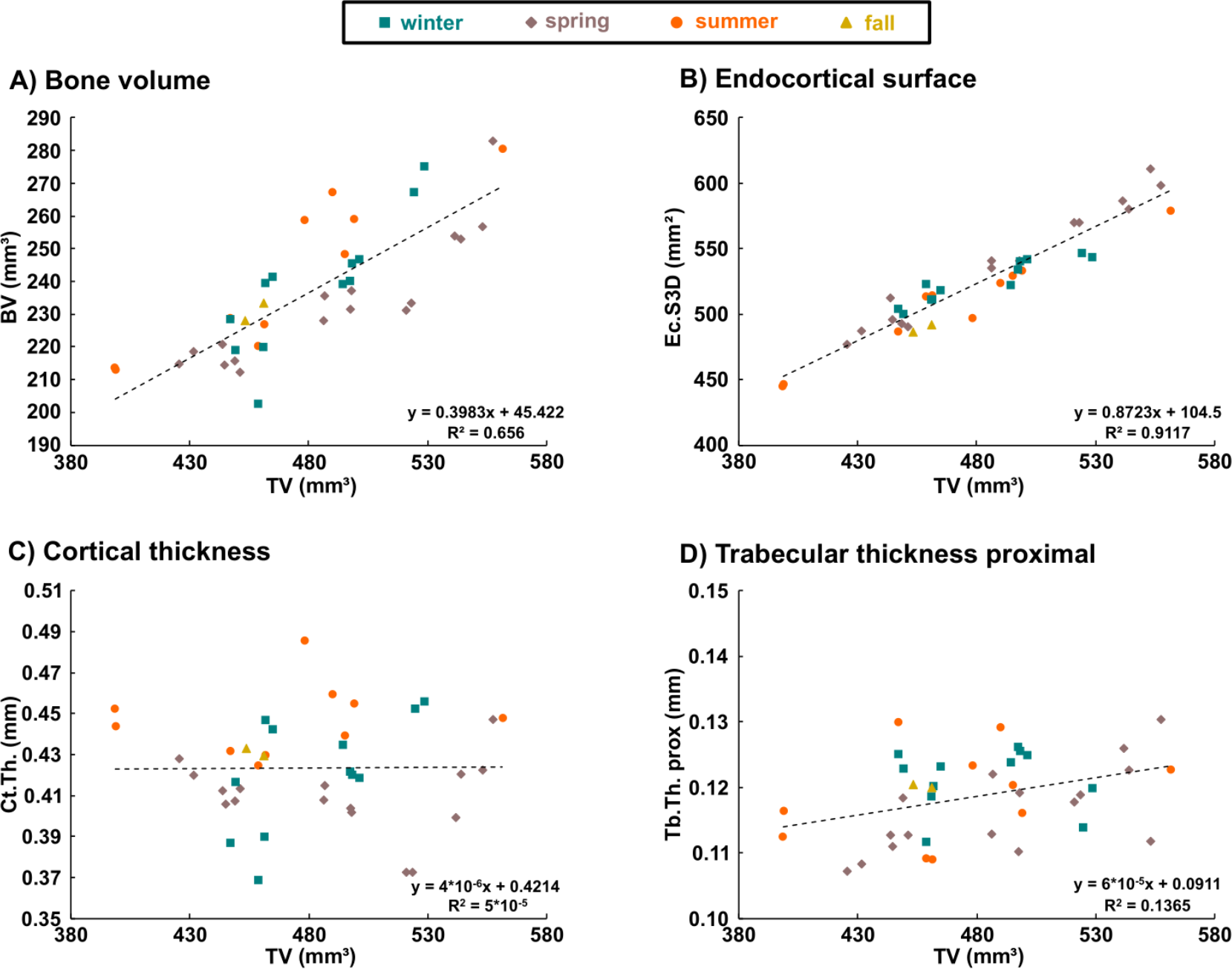
Multivariate linear regression of bone microstructure parameters onto total volume (TV) as a proxy for size. A. Average cortical thickness (Ct.Th.). B. Bone volume (BV). C. Endocortical surface (Ec.S3D). D. Mean trabecular thickness (Tb.Th. prox).

Bone microstructure as a whole (residual bone volume, cortical thickness, residual endocortical surface and residual trabecular thickness) is highly significantly different in estival vs hibernal squirrels (Pillai trace=0.3214, F(4, 35)=4.144, p=0.0075). Bone microstructure is also significantly influenced by season (Pillai trace=0.5067, F(8, 66)=2.799, p=0.0099). The post-hoc Hotelling’s T^2^ tests show that this significance is caused by a significant difference between spring and summer (p=0.0076, D^2^=3.4579). Season has a highly significant effect on residual endocortical surface, cortical thickness and residual bone volume (*Table 2* and *Figure 3*). According to the Tukey post-hoc tests, this is caused by highly Seasonal fluctuations in bone microstructure van Heteren et al. significant differences between spring and summer, and a significant difference between winter and summer in the case of cortical thickness. Additionally, the seasons significantly affect residual trabecular thickness (*Table 2* and *Figure 3*). According to the post-hoc Tukey tests, spring and winter are significantly different from each other. The seasons explain between 16% and 35% of the variance (*Table 2*) and the trendline explains between 14% and 33% of the variation (*Figure 3*).

**Table 2:**
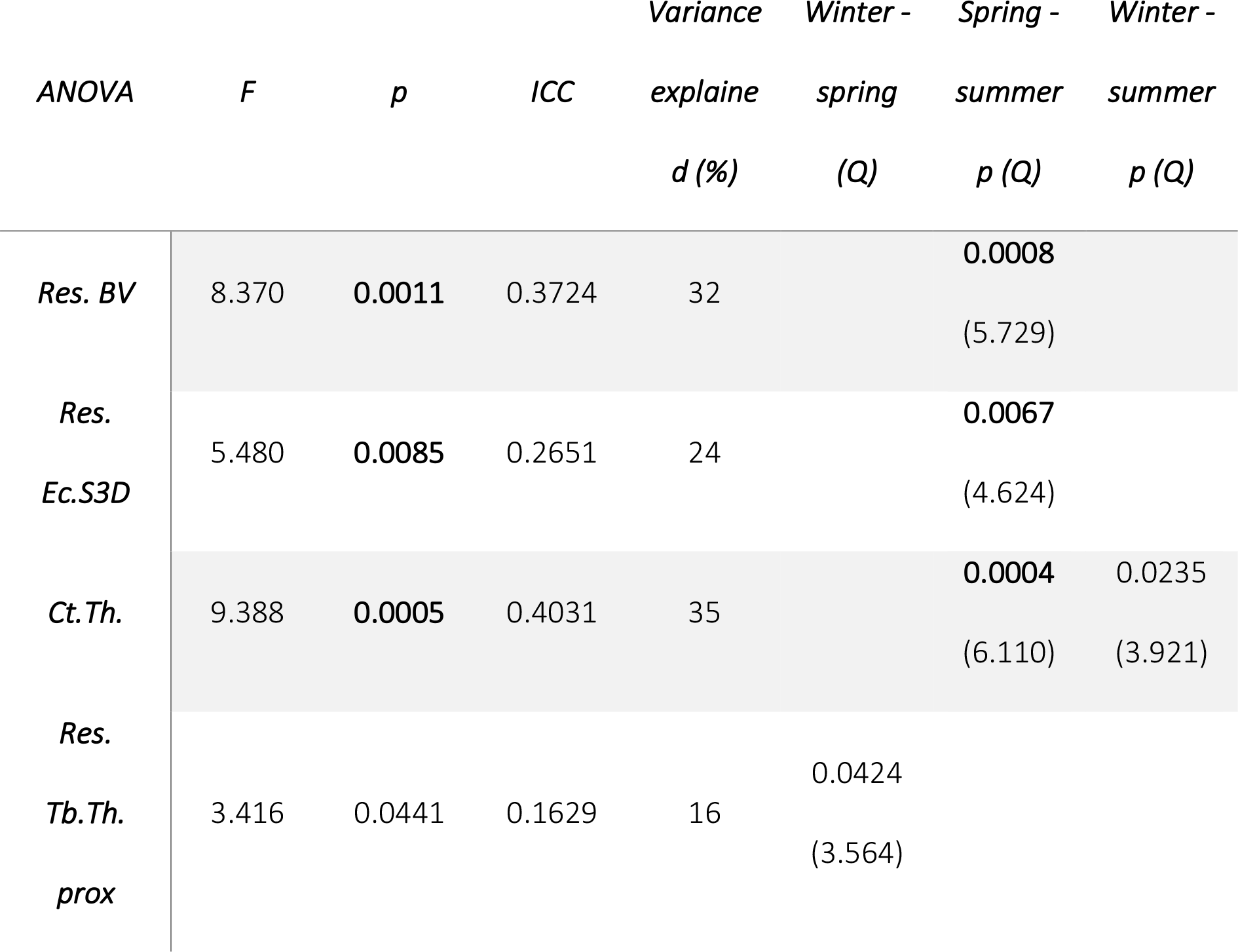
Results of the ANOVAs per season for residual bone volume (Res.BV), residual endocortical surface (Res.Ec.S3D), cortical thickness (Ct.Th.) and proximal trabecular thickness (Res.Tb.Th. prox) as well as the intraclass correlation coefficient (ICC) and the post-hoc Tukey tests. For the post-hoc tests, only significant values are provided. Highly significant values are indicated in bold font. For all F values, the between group degrees of freedom are 2 and the within group degrees of freedom are 35.

**Figure 3:**
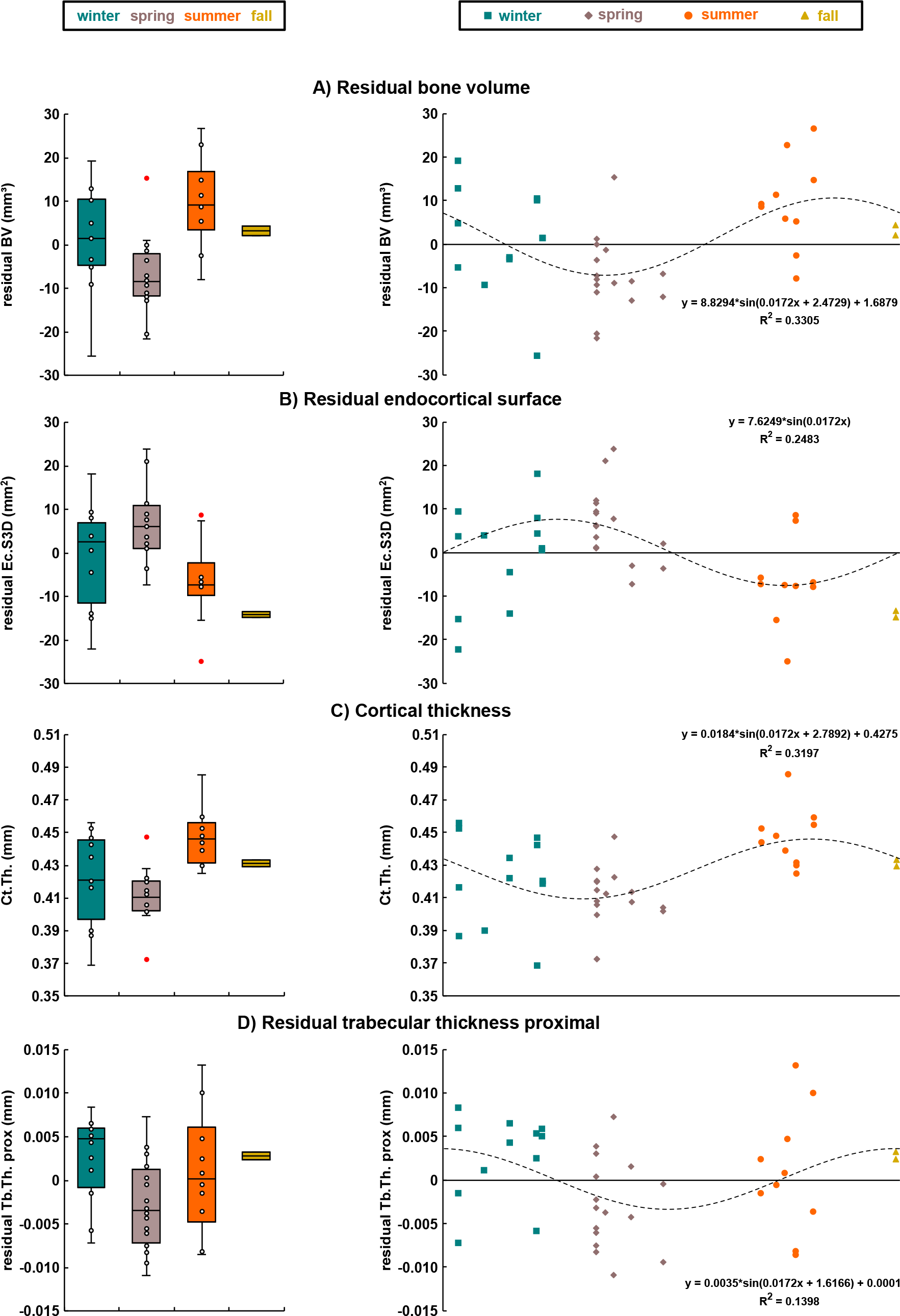
Boxplots (left) and scatterplots with sinus trendlines (right) of bone microstructure parameters. In the boxplots, the dots represent the datapoints, the length of the box is the interquartile range, the horizontal line is the sample median, and the whiskers extend to the minimum and maximum values, except for datapoints that are outside 1.5 times the interquartile range above the upper quartile or below the lower quartile, which are considered outliers and indicated in red rather than white. A. Average cortical thickness (Ct.Th.). B. Residual bone volume (Res.BV). C. Residual endocortical surface (Res.Ec.S3D). D. Residual mean trabecular thickness (Res.Tb.Th. prox).

## Discussion

Intraspecific variation in bone microstructure is rarely studied and, generally, humans are the focal taxon (e.g., Saers *et al*. 2018; Vom Scheidt *et al*. 2019). Most studies, however, focus on interspecific comparisons (e.g., Meier *et al*. 2013; Ryan & Shaw 2013; Mielke *et al*. 2018). In the present study, the intraspecific bone microstructure is analysed for a single subspecies of red squirrel (*S. vulgaris fuscoater*) from Bavaria (Germany) from a 11-year period at the beginning of the 20^th^ century, implying minimal geographical or temporal influences on the data. The squirrels used in this study are collection specimens from more than 100 years ago. Individual information on activity levels, breeding status, feeding behaviour or local environment is not available. Nevertheless, this is an important source of information. It does not require the sacrifice of additional animals, whether from the wild or from laboratory setting, both associated with their own ethical issues.

The effect of season on bone turnover is controversial (Rico *et al*. 1994; Woitge *et al*. 2000; Patel *et al*. 2001; Blumsohn *et al*. 2003; Seibel *et al*. 2004; Seibel 2005) and a previous study on microstructure parameters in sheep did not find any significant differences between the seasons (Arens *et al*. 2007). This study aims to uncover how the bone physiology of a non-human mammal responds to seasonality. The Central European squirrel (*S. vulgaris fuscoater*) was used as a model study system.

Significant differences were found between squirrels that were collected in the estival semiyear versus those that were collected in the hibernal semiyear. Subsequent analyses, with the year divided into four seasons, showed that this was mainly caused by a difference in the bone parameters in spring versus fall, with summer and winter as intermediate stages (*Figure 3*).

Almost 35% of the variance in bone volume was found to be related to season *(Table 2*). Since bone volume is comprised of the entire bone, both cortical and spongious, it is important to analyse further parameters to determine which aspects of the bone’s micromorphology might be responsible for such differences and how we can interpret those in terms of functional or eco-morphology.

In fall and winter the trabeculae are the thickest, in spring they are the thinnest, whereas squirrels display intermediate trabecular thicknesses in summer (*Figure 3*). Hazelnuts, especially, are very high in calcium (Łoźna *et al*. 2020; NutritionValue.org 2023) and primarily available at end of August and in September (Gurnell 1993), whereas acorns are particularly high in phosphorus (NutritionValue.org 2023) and are available from September to November with a noteworthy peak in the middle of November (Gurnell 1993). Their availability might allow for the thickening of the trabeculae over summer and fall.

Seasonal changes in cortical thickness seem to be shifted to earlier in the year relative to changes in trabecular thickness. Cortical thickness is higher in summer and fall than in winter and spring in squirrels (*Figure 3*). The seasonal pattern is the strongest in cortical thickness values; 35% of the variance is explained by season. Based on the available data, the mechanism behind the seasonal changes in cortical thickness remain unclear.

Endocortical surface is highest in spring and lowest in autumn, and season explains almost 25% of the variance in this parameter (*Table 2*). It shows a reversed pattern to bone cortical thickness, and they are essentially two sides to the same coin. When the periosteal surface stays the same throughout the seasons, but the endocortical surface increases or decreases, the cortical thickness also decreases or increases respectively. The same pattern is also observed in human smokers who have a smaller cortical thickness and a larger endocortical surface (Lorentzon *et al*. 2007). As such it would be plausible that a similar mechanism might be at play causing seasonal variations.

Summarising, this study showed that the cortical thickness increases highly significantly between spring and summer (*Table 2*) and decreases again in winter (*Figure 3*). Furthermore, trabecular thickness decreases from winter to spring (*Table 2*) and increases from spring to summer to fall (*Figure 3*). Both the categorical seasons as well as the gradual trendline, which is perhaps more in line with a squirrel-like perception of the environment explain close to one thirds of the variance.

Tree food availability for squirrels is highest in autumn, lowest in spring and intermediate in summer whereas winter was not assessed, (Reher *et al*. 2016), but is likely to be intermediate as well. Red squirrels depend on a variety of scatter-hoarded food types, when seeds are scarce in spring (Krauze-Gryz & Gryz 2015). Not all animals are equally successful and those that retrieve more cached tree seeds are more likely to survive the spring breeding season (Wauters *et al*. 1995).

Red squirrels are known to predate on eggs, juvenile birds (Lurz *et al*. 2005), which, in Europe, tend to be available between March and June (Lack 1950), and other animals (Moller 1983a). In grey squirrels, % animal matter in the stomach is highest in spring and summer (Moller 1983a) and red squirrels are likely to show similar behaviour. Until now, it was not clear whether squirrels predate on other animals to obtain proteins (i.e., meat) or minerals like (i.e., calcium and/or phosphorus) (Callahan 1993), although American red squirrels (*Tamiasciurus hudsonicus*) (Leech 1977) and Eastern fox squirrels (*Sciurus niger*) (Callahan 1993) have been reported to eat bones, suggesting the latter for those species. Grey squirrels have been reported to strip tree bark and eat the phloem (Nichols *et al*. 2016). This was thought this counteracted to a seasonal calcium deficiency (Nichols *et al*. 2016), but more recent research has shown that grey squirrels are unable to utilise calcium oxalate, the form in which calcium is available in phloem (Nichols *et al*. 2018). Red squirrels, being closely related to grey squirrels, are also unlikely to be able to utilise the calcium from phloem and must obtain it elsewhere. The present study suggests that Eurasian red squirrels might predate to replenish their calcium and/or phosphorus, in addition to eating hazelnuts and acorns, so their bones can recover. The lower cortical thickness detected in the hibernal semiyear samples and the delayed lower trabecular thickness in spring and summer could potentially be explained by the role of bone as a calcium reservoir. Calcium is messenger which couples intracellular responses to extracellular signals, for example the activation of muscle contraction (Awumey & Bukoski 2006). Since squirrels do not hibernate, calcium must be used throughout winter. In the case of low calcium availability in the diet, the skeleton might possibly be used for bone resorption (Heaney 2006). This might result in the observed decrease in trabecular thickness over the winter months.

There are also alternative explanations. Disuse of bones, such as in hibernating mammals, also leads to bone loss, because bone formation and bone resorption become unbalanced (McGee-Lawrence *et al*. 2008).The red squirrel is not a hibernating species, nevertheless physical activity is reduced in winter months (Tonkin 1983). Whether this also contributes to fluctuations in bone parameters cannot presently be excluded and would require experimental research.

Bone homeostasis is maintained by osteoclastic-osteoblastic activity (Guo *et al*. 2018), as well as osteocytic osteolysis (Tsourdi *et al*. 2018). The present study does not provide enough information to assess which of these processes plays the most important role, but both osteoclastic-osteoblastic activity and osteocytic osteolysis are affected by vitamin D (Lanske *et al*. 2014; Takahashi *et al*. 2014; van Driel & van Leeuwen 2014). Vitamin D is produced by the body under the influence of sun light and cannot be taken up through food. Since the days are shorter and red squirrel activity is reduced in winter (Tonkin 1983), red squirrels would be expected to produce less vitamin D in the colder months. The influence of vitamin D on bone mineralisation is complex and it seems to stimulate osteoblast mineralisation in humans, but the effect on mineralisation in murines (Old World rats and mice) is not uniform (van Driel & van Leeuwen 2017) and vitamin D can both positively and negatively regulate osteoblasts in rats (Owen *et al*. 1991). Since red squirrels are rodents too, their physiological response to fluctuations in vitamin D availability cannot be predicted and will have to be assess experimentally.

## Acknowledgements

The authors would like to thank Stefan Filser (Bavarian State Collection of Zoology) and Thijs Heesters for their assistance, Alexander Floroni (Ludwig-Maximilans-Universität München) for contributions to the artwork, Yves Maris (University of Antwerp) for IT support, and Mathieu Gendron, Nicolas Piché and Emimal Jabason (Object Research Systems) for methodological advice and help. Funding was provided by the Bayerische Staatsministerium through the Pakt für Forschung und Innovation as an SNSB Innovativ grant.

## Supplementary material

All supplementary information can be found in a collection on Figshare at https://doi.org/10.6084/m9.figshare.c.6435755. Raw data (Suppl. Info. 1) are available at https://doi.org/10.6084/m9.figshare.22121447. Alternative statistics using specimen means instead of bilateral data can be found in Suppl. Info. 2 here: https://doi.org/10.6084/m9.figshare.24716520. Code for the reconstruction of the CT scans (van Aarle *et al*. 2015; van Aarle *et al*. 2016) is not novel, but is provided at https://doi.org/10.6084/m9.figshare.22121441 for ease of use as Suppl. Info 3. Novel code for isolating bone tissue in Dragonfly (Suppl. Info. 4) is available at https://doi.org/10.6084/m9.figshare.22121456.

